# Chemoprophylaxis vaccination with a *Plasmodium* liver stage autophagy mutant affords enhanced and long-lasting protection

**DOI:** 10.1101/2020.10.23.352799

**Authors:** Tejram Sahu, Ella J. Gehrke, Yevel Flores-Garcia, Godfree Mlambo, Julia D. Romano, Isabelle Coppens

**Affiliations:** Department of Molecular Microbiology and Immunology, Johns Hopkins Malaria Research Institute, Johns Hopkins University Bloomberg School of Public Health, Baltimore 21205, MD, USA

## Abstract

Genetically-attenuated sporozoite vaccines can elicit long-lasting protection against malaria but pose risks of breakthrough infection. Chemoprophylaxis vaccination (CVac) has proven to be the most effective vaccine strategy against malaria. Though CVac with WT sporozoites confers better immunity, the overhanging threat of drug resistance limits its use as a vaccine. Here, we demonstrate that a liver stage-specific mutant of *Plasmodium berghei* when used as a vaccine under a CVac regimen provides superior long-lasting protection, in both inbred and outbred mice, as compared to WT-CVac. Uniquely, the protection elicited by this mutant is predominantly dependent on a CD8^+^T-cell response through an IFN-γ-independent mechanism and is associated with a stable population of antigen-experienced CD8^+^T cells. Jointly, our findings support the benefit of liver stage mutants as vaccines over WT, under a CVac protocol. This vaccination strategy is also a powerful model to study the mechanisms of protective immunity and discover new protective antigens.

## Introduction

Malaria caused by *Plasmodium* parasites remains an infectious disease of major public health importance. Malaria vaccine development is a continuously evolving field of research, which aims towards a vaccine with long-lasting stage, strain and species transcending sterile immunity. Such an ideal vaccine is more essential than ever in the face of increasing resistance to frontline drugs by malaria parasites. The recently licensed malaria subunit vaccine (RTS,S) using genes from the repeat and T-cell epitope in the circumsporozoite protein of *Plasmodium falciparum* (PfCSP) conjugated to a hepatitis B virus surface antigen in addition to ASO-type adjuvants, induces humoral and cellular immunity^1,2^. RTS,S/AS0 vaccines reduce episodes of severe malaria and delay the time to new infection but do not offer a durable, high level efficacy (> 50%). Vaccination using whole sporozoites (*Plasmodium* forms in mosquito salivary glands), though logistically challenging, bears much promise due to exposure of a large repertoire of immunogens to the immune system^3–9^. Among whole sporozoite vaccines (WSV), Radiation-Attenuated Sporozoites (RAS) has been considered the “gold standard” model for induction of full and sterilizing immunity against malaria^5,10^.

Afterwards, a reverse vaccinology approach based on *Plasmodium* genomes^11^, bioinformatics tools predicting antigens and/or essential proteins, and improvements in transfection methods for *Plasmodium* genetic manipulation^12–14^ has led to the generation of several parasite stage-specific knock-outs exploitable as Genetically Attenuated Parasite (GAP) vaccines^7,15,16^. GAP were designed to overcome the uncertainty associated with random DNA breaks in RAS and minimize the risks associated with radiation, including the production of free radicals. However, in almost all cases, GAP were limited by their breakthrough to blood stage, rendering them unsuitable for clinical application (reviewed in^17^). In addition, while RAS arrest at the onset of liver stage development and express primarily sporozoite-derived antigens, GAP can be designed with arrest either toward the mid or end of liver stage development. The selection of the candidate gene for manipulation to engineer a GAP-vaccine is critical as the GAP must be safe (i.e. giving a complete attenuation in the liver) and potent (i.e. generating a strong protective immunity). A GAP with a late liver stage arresting phenotype would induce stronger and broader protective immune responses.

*Plasmodium* parasites undergo continuous cellular renovation to adapt to various environments in their vertebrate host and insect vector. We previously showed that during the conversion of sporozoites to liver forms in hepatocytes, the malaria parasite upregulates its autophagy machinery to eliminate sporozoite organelles that are not essential for parasite replication in the liver^18,19^. Our studies have identified *Plasmodium* ATG8 as an important autophagy protein involved in parasite metamorphosis in the liver. In the attempt to dysregulate parasite autophagic functions, we engineered a mutant of a *Plasmodium berghei* line with a modified ATG8 3’UTR that overexpresses ATG8 by ~2-fold (PbATG8-OE)^18^. PbATG8-OE parasites are competent to initiate a liver infection but suffer a mid-liver stage developmental defect, with a delay in expulsion of sporozoite organelles, such as micronemes involved in motility and invasion.

Recently, infection with WT sporozoites under a chemoprophylaxis treatment (commonly with chloroquine) named CVac is exploited as a vaccination approach. CVac has been proven to provide long lasting sterilizing protection with 2-log less sporozoite antigens as compared to RAS^6,8,20^. However, under a CVac regimen, the development of resistance to a prophylactic drug would be a limiting factor for its wide field application. In this study, we monitored in more detail, the development of PbATG8-OE parasites in cultured hepatocytes and mouse liver over time. We also explored the potential of this mutant to elicit a protective immune response in immunologically relevant mouse models, under a CVac regimen. We systematically evaluated the mechanisms of immune protection induced by the mutant, in comparison with parental parasites, with the premise to shed light on the priming and sustenance of immune responses against pre-erythrocytic parasites and the potential use of GAP under a CVac regimen as a vaccination strategy for malaria control.

## Methods

### Ethics approval

All animal procedures were approved by the Institutional Animal Care and Use Committee of the Johns Hopkins University following the NIH guidelines for animal housing and care. Infected mice were euthanized when the parasitemia reached 20% or when they presented signs of severe discomfort and/or morbidity.

### Mice, mosquitoes, parasites, and cell lines

All purchased mice were 6-week-old females and housed in the Johns Hopkins Bloomberg School of Public Health animal care unit. Swiss-Webster mice were bought from Enviligo Harlan Laboratories Inc. (Frederik, MD). Balb/c mice, C57BL/6J mice, IFN-γ-KO mice (Stock# 002287) and IFN-γR-KO mice (Stock# 003288) were purchased from Jackson Laboratory (Bar Harbor, ME). For all our experiments, mice were fed *ad libitum*. Protocols for mice immunization are detailed in the figures and described in the figure legends. *Anopheles stephensi* mosquitoes, infected with appropriate strain of *Plasmodium*, were provided by the JHMRI Parasitology Core Facility. Parasite strains included: WT-FLP from *Plasmodium berghei* (Pb-ANKA), PbATG8-OE^18^, *P. berghei*-mCherry^21^ (gift from Photini Sinnis, Johns Hopkins University), *P. berghei* sporozoites expressing PfCSP^22^ generously provided as air-dried sporozoites from Fidel Zavala (Johns Hopkins University) and Pb-GFP-Luc^13^ (MRA-868) sporozoites obtained through the JHMRI Parasite Core Facility. Cryopreserved parasites were passaged once through Swiss-Webster mice before infection in experimental animals. Mouse Hepa1-6 cells (ATCC CRL-1830) used to monitor infection *in vitro* were obtained from ATCC (Gaithersburg, MD). Cells were grown as monolayers at 37°C in an atmosphere of 5% CO_2_ in α-MEM supplemented with 10% FBS, 2 mM L-glutamine and penicillin/streptomycin (100 U/ml).

### Parasitemia and liver stage parasite burden estimation

To study malaria blood stage, mice were infected intravenously (i.v.) with appropriate number of *Plasmodium*-infected red blood cells (iRBC) from a donor mouse, and the ensuing parasitemia was assessed microscopically by iRBC enumeration on giemsa-stained thin blood smears (at least 25 fields from 200 iRBC/field for each smear) or by PCR for Pb 18S rRNA gene. Negative blood stage infection after 2 weeks was further verified by transfer of 100 μl of blood from infected to naïve mice by tail i.v. injection and monitoring blood stage infection for 15 days on thin blood smears. For sporozoite infection, parasites were cycled between Swiss-Webster mice and *A. stephensi* mosquitoes. To collect sporozoites, salivary glands dissected from infected mosquitoes on D21-D25 post-blood feeding were disrupted by passage 20 times through a 27.5G needle fitted to a 1 ml syringe. Mice were infected by i.v. injection of sporozoites through tail vein, and blood patency was monitored beginning D3 on blood smears. To assess liver stage parasite burden after sporozoite infection of mice, livers were isolated 40 h p.i. from animals to extract total RNA with TRizol (Thermofisher Scientific, Waltham, MA) using RNeasy mini kit (Qiagen Inc., Germantown, MD) as described^23^. DNA was removed from total RNA preparations using On-Column DNase I Digestion Set (Sigma Aldrich, St. Louis, MO). cDNA was synthesized from 1 μg of total RNA using Super Script III First Strand cDNA Synthesis Kit (Invitrogen, Carlsbad, CA) and random hexamer. A Standard curve quantitative RT-PCR was performed in a 25 μl volume using ABI Power SYBR Green Master Mix (Thermofisher Scientific), with 1:12.5 dilution of cDNA and 0.25 μM of *P. berghei* 18S rRNA primers (forward-5’-AAGCATTAAATAAAGCGAATACATCCTTAC-3’ and reverse-5’-GGAGATTGGTTTTGACGTTTATGTG-3’) or mouse β-actin primers (forward-5’-GGCTGTATTCCCCTCCATCG-3’ and reverse-5’-CCAGTTGGTAACAATGCCATGT-3’). Standard curves were generated using known gene copy numbers from 10^3^ to 10^8^ of Pb-18S rRNA and mouse β-actin diluted in 2 μl of nuclease-free water. PCR reactions were run on ABI 7500 Fast Real-Time PCR System, using the following thermal cycling conditions: 95° C for 15 min, 40 cycles with 95° C for 20 s; 60° C for 30 s, and 72° C for 50s. Rhodamine-X (ROX) was used as passive reference dye for normalization of the fluorescence intensity generated by qPCR using SYBR Green method.

### Quantification of MSP4/5 transcripts during late liver stage

For MSP4/5 transcript quantification, mouse livers were harvested after perfusion with 10 ml of RNAse-free PBS (Thermofisher Scientific), through hepatic portal vein 65 h post-sporozoite injection. Total RNA was isolated from infected liver, and cDNA was synthesized as described above. Transcript abundance was assessed using a qPCR standard curve method as described^24^, except that genomic DNA was replaced by known copy numbers of PCR-amplified targets as standards in our assays. Real-Time qPCR was performed using the primers forward-5’-GAAAGCCGTAAATTACTTATCACTG-3’ and reverse-5’-CCCTCATTTTGATTCGAACTAGTTG-3’ for MSP4/5 and forward-5’-TGCAGCAGATAATCAAACTC-3’ and reverse-5’-ACTTCAATTTGTGGAACACC-3’ for Hsp70, cDNA from the infected liver preparations and the SYBR Green Master Mix. The PCR conditions were: 95° C for 10 min, 40 cycles of 95° C for 10 s, 42° C for 20 s and 60° C for 40 s. Relative abundance of MSP4/5 was calculated by comparing its abundance with that of the Hsp70 transcripts.

### Mosquito bite challenge

For each mosquito bite challenge, mice immunized with WT-FLP-CVac or PbATG8-OE-CVac and naïve mice were anesthetized and placed on the top of a cup containing 10-12 Pb-ANKA-infected female *Anopheles* mosquitoes. Mosquitoes were allowed to feed for 30 min on the mice prior to dissection. Mosquitoes were then monitored by visual observation for the presence of blood in the gut and sporozoites within salivary glands. Both a midgut engorged with blood and salivary glands containing sporozoites were accounted for an infectious mosquito bite. Development of blood infection in mice was monitored 3 days post-mosquito bites until D15 on thin blood smears.

### In vivo T-cell depletion

For selective T-cell depletion from PbATG8-OE-CVac-immunized mice, 500 μg of rat anti-CD8 (clone 2.43; BioXCell, West Lebanon, NH), rat anti-CD4 (clone GK 1.5, BioXCell) monoclonal antibodies or isotype control (LTF-2, BioXCell) were i.p. injected into immunized mice one day prior to and on the day of the challenge^25^.

### Quantitative *Plasmodium* Sporozoite Neutralization Assay (SNA)

To determine the anti-sporozoite immunity, SNA was performed as described^26^ with minor modifications. Pb-ANKA (20,000 sporozoites) freshly isolated from salivary glands were with diluted serum (1:6 in PBS) collected from either immunized or naïve mice in a 30 μl volume on ice for 45 min. Serum-exposed sporozoites were intradermally (a clinically relevant route of infection) injected into naïve mice to monitor development of blood infection by blood smearing starting from D3 post-sporozoite injection until D15.

### Immunophenotyping of Parasite-Specific CD8^+^ T Cells

Peripheral blood of WT-FLP-CVac and PbATG8-OE-CVac-immunized mice were immunostained with anti-CD8a (53-6.7; eBioscience, San Diego, CA) and anti-CD11a (M17/4; eBioscience) to identify CD8α^lo^CD11a^hi^ T cells from the total CD8^+^ T cell population^7,27^. This subset of T cells were analyzed by flow cytometry using Attune Next Flow Cytometer (Thermofisher Scientific) and data analyzed using FlowJo Software (Version 10.6.1) from Tree Star Inc. (Ashland, OR). Mouse blood was withdrawn prior to immunization to establish basal level of peripheral CD8α^lo^CD11a^hi^ T cells, and post-priming, post-boost and 3-month post-boost to monitor the changes in the number of these CD8^+^ T cell populations.

### Blood stage challenge assay

To assess cross-stage protection, mice were immunized with 2 or 3×20,000 PbATG8-OE-CVac or WT-FLP-CVac parasites. Both immunized and naïve mice were i.v. infected with 10^5^ iRBC at D21 or D80, and blood stage infection was monitored on thin blood smears.

### Indirect Immunofluorescence assay (IFA) and parasite imaging

<u>Sporozoites:</u> IFA were performed on sporozoites isolated from salivary glands, mixed blood stages and exoerythrocytic forms (EEF) as described elsewhere^18,28^, with few modifications. To evaluate the antibody response generated against sporozoite antigens following immunization, sporozoites were incubated for 45 min in a poly-L-lysin-coated 8-well Lab-Tek chambered slide (LabTek Inc., Grand Rapids, MI), fixed with 4% PFA+0.02% Glutaraldehyde in PBS for 15 min at room temperature, washed twice with PBS, blocked with 3% BSA in PBS overnight at 8°C, and exposed to sera diluted at 1:400 from immunized or naïve mice. After washing four times with PBS, sporozoites were incubated with Alexa-488 conjugated anti-mouse IgG (Thermofischer Scientific) for 2 h at a dilution of 1:1,000 in 3% BSA/PBS. Slides were then washed three times (15 minutes each) with PBS to remove unbound antibodies and stained with DAPI (1 μg/ml). <u>Blood forms:</u> *P. berghei*-mixed infected blood was collected by tail snip. Fixation and immunostaining protocol were adapted from^29^ with minor modifications. Briefly, iRBC in eppendorf tubes were washed with PBS by centrifugation at 800x*g* for 5 min at room temperature, fixed with 4% PFA+0.007% glutaraldehyde (EM grade; Electron Microscopy Sciences, Hatfield, PA) in PBS for 30 min at room temperature, washed with PBS, permeabilized with 0.1% Triton X-100 in PBS for 10 min, washed with PBS and treated with 0.1 mg/ml of NaBH_4_ at room temperature for 10 min to remove free aldehydes. Cells were then washed again once with PBS, blocked overnight with 3% BSA in PBS at 8° C, incubated with sera diluted at 1:400 from immunized mice overnight at 8° C, washed 3 times with PBS and then incubated with 1:1,000 dilution of Alexa-488 conjugated anti-mouse IgG for 1 h at room temperature. iRBC preparations were then spotted onto a coverslip pretreated with polyethyleneimine (PEI), and parasite-containing coverslips were washed with PBS and mounted with ProLong Gold antifade reagent (Thermofisher Scientific) with DAPI. <u>Exoerythrocytic forms (EEF):</u> confluent Hepa1-6 cells seeded in a 24-well plate were infected with 5,000 freshly dissected sporozoites. After centrifugation of the plate at 400x*g* for 5 min for sporozoites contacting mammalian cells, monolayers were incubated at 37°C for 3 h before washing with PBS to remove extracellular (noninvading) sporozoites. Infected cells were washed, fixed and immunostained at different time points as described^18,28^. Fixed samples were viewed with either a Nikon Eclipse 90i fluorescence microscope equipped with an oil-immersion Nikon plan Apo 100x/NA 1.4 objective, or a Nikon plan Fluor 60x/NA 0.75 objective and a Hamamatsu GRCA-ER camera (Hamamatsu Photonics, Hamamatsu, Japan) or a Zeiss AxioImager M2 fluorescence microscope equipped with an oil-immersion Zeiss plan Apo 100x/NA 1.4 objective and a Hamamatsu ORCA-R2 camera. Optical z-sections with 0.2μm spacing were acquired using Volocity 6.3.1 software acquisition module (Perkin Elmer, Waltham, MA).

### In vitro assessment of *Plasmodium* liver stage development

To assess liver stage development, 10^5^ Hepa1-6 cells were grown overnight on coverslips in a 24-well plate as described above. Cells were then infected with 20,000 sporozoites per well, fixed and stained for DAPI and PbHsp70 by IFA. Number of infected cells per well were counted by fluorescence microscope and images with 0.2 μm optical *z*-section were acquired using a Zeiss AxioImager and a 63x oil-immersion objective. EEF PV volume were quantified by Volocity software using Hsp70 expression by the parasite. To measure nucleic acid in developing EEF, images were acquired at the same light intensity (20%) and exposure time (60-80 ms, depending on the time point after infection) between strains and the DAPI signal was measured using Volocity software. DAPI intensity signal was expressed as sum of total DAPI fluorescence intensity in arbitrary units.

### Merosome isolation and counting

Hepa1-6 cells were infected with 20,000 sporozoites per well and were grown for 65 h as described above. Culture supernatant were collected at 65 h, and floating merosomes and detached cells were spun down at 2,000*g* for 5 min at 4° C as described^24^. Pellet containing merosomes and cells were further washed twice with cold PBS by centrifugation and resuspended in 100 μl of PBS from which 10 μl was used for counting using C-Chip disposable hemocytometer (INCYTO, SKC Inc, Covington, GA).

### Transmission electron microscopy

For ultrastructural observations of blood forms by thin-section transmission electron microscopy (EM), iRBC were fixed in 2.5% glutaraldehyde in 0.1 mM sodium cacodylate (EMS) and processed as described^19^. Ultrathin sections of infected cells were stained before examination with a Philips CM120 EM (Eindhoven, the Netherlands) under 80 kV.

### Statistical Analysis

Data were analyzed using Graph Pad Prism8 software. Specific tests performed with corresponding statistical significance are specified in each figure legend.

## Results

### PbATG8-OE parasites have significant delay in blood infection after sporozoite injection into mice

In our earlier study, we reported that *Plasmodium* parasites express ATG8 throughout the lifecycle, but selectively upregulate their ATG8 ubiquitin-like conjugation system during liver stage, as demonstrated for *P. berghei* in hepacytes^18,19^. The parasite autophagy machinery is a prerequisite for sporozoite differentiation into liver forms and is involved in the elimination of sporozoite organelles useless for replication. We showed that a *P. berghei* strain engineered for PbATG8 overexpression (PbATG8-OE) can invade hepatocytes *in vitro* but poorly develops, as compared to parental parasites (WT-FLP)^18^. A preliminary investigation using 3 outbred mice infected with 2 doses of sporozoites (5,000 and 50,000) reveals that PbATG8-OE parasites were unable to form infectious exoerythrocytic merozoites^18^. We wanted to expand this observation by infecting two different strains of mice (outbred Swiss-Webster and inbred BALB/c mice, with 5 mice per group in duplicate assay) with 10,000 freshly dissected sporozoites, to determine the amount of time spent by the mutant in the liver before initiating blood stage infection (Fig. 1a). Blood infection in mice monitored on blood smears shows that the prepatent periods (i.e., the time until the first day of parasite detection in erythrocytes) were significantly longer for PbATG8-OE parasites (7 to 12 days vs. 3 to 5 days in WT-FLP parasites. Moreover, a subset of mice (20% of Swiss-Webster and 60% of BALB/c mice) remained blood smear-negative up to 15-day post-sporozoite infection. These additional sets of data suggest that PbATG8-OE parasites stay longer in the liver before emerging into the blood, confirming growth delay in hepatocytes.

**Figure 1.**
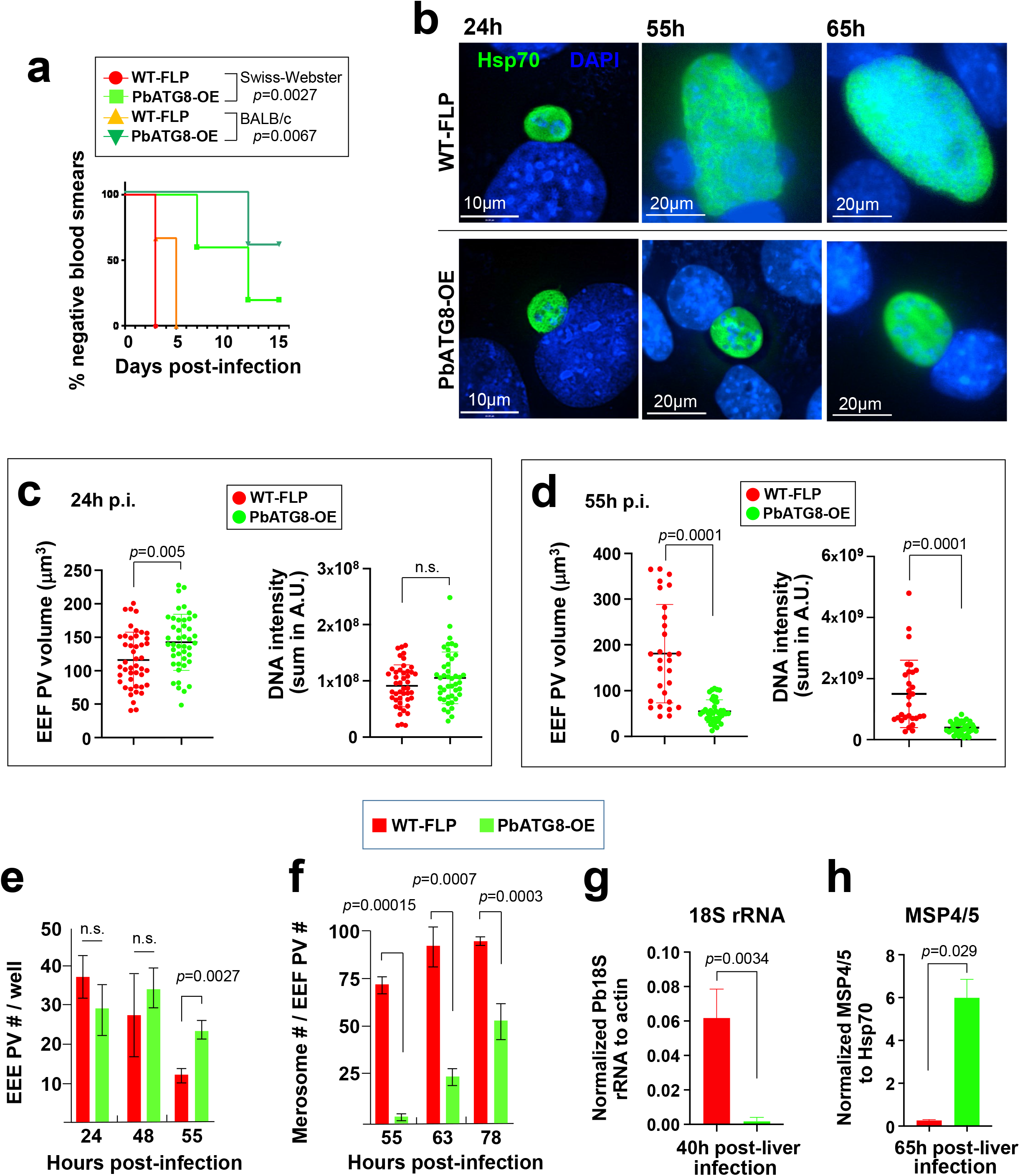
PbATG8-OE parasites have severe liver stage developmental defects. **a.** Infectivity *in vivo*. Kaplan-Meier survival plot showing days to blood stage infection after infection with 10^4^ sporozoites from WT-FLP (red circles and orange triangles) or PbATG8-OE (light green squares and dark green inverted triangles) parasites in outbred Swiss-Webster and inbred Balb/c mice (n=5 in each group; *p* values determined by log-rank Mantel Cox test). **b-f.** Development in vitro. IFA showing representative images of developing EEF of WT-FLP and PbATG8-OE parasites in Hepa1-6 cells at 24, 55 and 65 h p.i. in **b**. Parasites were stained with antibodies against PbHSP70. DAPI in blue. Quantification of EEF size (volume in μm^3^) and nucleic acid content for schizogony (measured by DAPI intensity) at 24h (**c**) and 55 h p.i. (**d**) of WT-FLP or PbATG8-OE parasites in Hepa1-6 cells. Nucleic acid content was assessed as sum of total DAPI intensity and expressed in arbitrary Unit (A.U.). Data are means ± SD (n=45 EEF in **c** and n=29-39 in **d**; *p* values determined by Mann-Whitney test). Enumeration of EEF PV (**e**) and merosomes released into the culture supernatant (**f**) of WT-FLP and PbATG8-OE parasites over time, based on HSP70 fluorescence. Data are means ± SD (n=4; p values determined by Mann-Whitney test). **g-h.** Mice liver assays: Swiss-Webster mice were i.v. infected with 20,000 sporozoites from WT-FLP or ATG8-OE. In **g**: livers were harvested at 40 h p.i. to assess parasite burden by measuring parasite 18S rRNA copy number by RT-qPCR. Data were normalized with host β-actin copy number and expressed as a ratio of parasite/host. In **h**: livers were harvested at 65 h p.i. after perfusion (to remove blood stage parasite contamination) to assess schizont maturation based on relative abundance of *Plasmodium* MSP4/5 measured by RT-qPCR. MSP 4/5 copy number were estimated by standard curve qPCR and normalized to parasite HSP70 copy number. Data in **g** and **h** are means ± SD of 5 mice per group per time point. Statistical significance was determined by Mann-Whitney test.

### PbATG8-OE parasites display growth defects, both in size and DNA replication from mid-liver stage

We next examined whether this observed delay in prepatent period of PbATG8-OE parasites is due to an abnormally slow, prolonged development of exoerythocytic forms (EEF) in hepatocytes or a release of only a few hepatic merozoites from cells, or a combination of both. We set up an in vitro infection system using Hepa1-6 cells infected with sporozoites to monitor the development of the mutant parasite at different time points, from 24 h post-infection (p.i.) to 78 h. Exoerythrocytic schizogony is characterized by the expansion of the parasite cytoplasmic mass and by multiple rounds of DNA replication that generate thousands of individualized nuclei^30^. We performed immunofluorescence assays (IFA) using antibodies against parasite Hsp70 to observe the parasitophorous vacuole (PV) size and DAPI staining to monitor parasite DNA content (Fig. 1b). The intensity of the Hsp70 and DAPI fluorescence signals was measured for quantitative comparisons between mutant and parental parasites (Fig. 1c and d). Results show that PbATG8-OE and WT-FLP EEF had a comparable development up to 24 h p.i., with the same PV size and DNA abundance. However, at 55 h p.i., the mutant had barely expanded in size, and exhibited strong defects in the extent of DNA replication. The end of schizogony is characterized by the compartmentalization of the cytoplasm containing organelles and a nucleus, leading to the formation of thousands of individualized hepatic merozoites^31^. Merozoites then exit host cells in the form of merosomes, i.e., hepatocyte-derived vesicles containing hundreds of parasites. At 65 h p.i., the vast majority of parental parasites had egressed from hepatocytes as expected while PbATG8-OE EEF were still intracellular (not shown). To document further schizogony and egress defects in the mutant, we enumerated the EEF per well seeded with hepatic cells during early (24 h p.i.), mid (48 h p.i.) and late (55 h p.i.) liver stage development (Fig. 1e). No significant difference in the number of EEF per well was observed between the two strains at 24 h and 48 h p.i., however, a significantly higher number of PbATG8-OE EEF were observed at 55 h p.i. Counting the number of free floating merosomes in the culture supernatant relative to EEF at 55 h, 63 h and 78 h p.i. confirms a productive formation of merosomes for parental parasites, with close to ~95% of liver forms being hepatic merozoites at 78 h p.i. (Fig. 1f). By that time, PbATG8-OE parasites exhibited a significant delay in merosome production, with about half of the parasite population unable to reach the merosomal stage, and thus presumably dead within the host cell. These observations reveal that PbATG8-OE parasites suffer from pronounced growth delays during the late stages of schizogony, resulting in a significant decrease in merosome formation and budding from cells.

These in vitro observations were then verified in infected animals in which parasite liver stage burden and maturation into merozoites in mouse liver were assessed by qPCR at 40 h (mid-liver stage development for *P. berghei* WT) and 65 h (post-liver development). Parasite load based on Pb18S rRNA copy number at 40 h post-sporozoite injection was significantly lower for PbATG8-OE parasites compared to parental parasites, confirming the slow, defective replication of the mutant in the liver (Fig. 1g). At 65 h, parasite burden and differentiation to late stage merozoites was performed based on relative abundance of MSP4/5 (a merozoite-specific plasma membrane marker) to Hsp70 in the perfused infected liver. As expected for the parental strain, almost no gene copies of MSP4/5 were detected, indicating egress from the liver. However, the copy number of MSP4/5 was significantly higher for PbATG8-OE parasites, revealing a long-lasting liver stage in mice infected by the mutant (Fig. 1h).

### PbATG8-OE parasites undergo normal asexual blood stage development

We next examined if the delayed blood prepatency observed with PbATG8-OE parasites could be due to blood stage defects, in addition to liver stage defects. The blood stage development of the mutant was monitored following mouse injection of RBC infected with PbATG8-OE parasites, in comparison to infected RBC (iRBC) with WT-FLP and WT-FRT (strain with the recombination construct of FRT sites but before the recombination event). No difference in parasitemia was observed between the 3 strains up to 4 days post-iRBC injection (Fig. 2a). The presence of the mutation introduced in the 3’UTR of *Atg8* (insertion of extra 60-bp sequence) in PbATG8-OE was verified by PCR using primers specifically designed to amplify bands in parental and PbATG8-OE but not in FRT parasites, with an expected size shift in PbATG8-OE blood forms (Fig. 2b). We further confirmed the presence of the mutation by sequencing of the 3’UTR of the *Atg8* gene. To monitor the stability of the genetic modification introduced in the PbATG8-OE strain, mutant parasites were cycled repeatedly (at least 5-times) through both mice and mosquitoes, and analysis of genomic DNA showed the continued presence of the 60-bp sequence, indicating no genetic reversion to WT. We also performed studies to inspect the ultrastructure of PbATG8-OE and WT-FLP blood forms, 5- and 18-day post-inoculation of sporozoites into mice. EM observations did not reveal any abnormalities in organellar composition and expansion in the mutant infecting RBC (Fig. 2c). Thus, the genetic modification introduced in the *Atg8* locus does not appear to affect blood stage growth of PbATG8-OE parasites.

**Figure 2.**
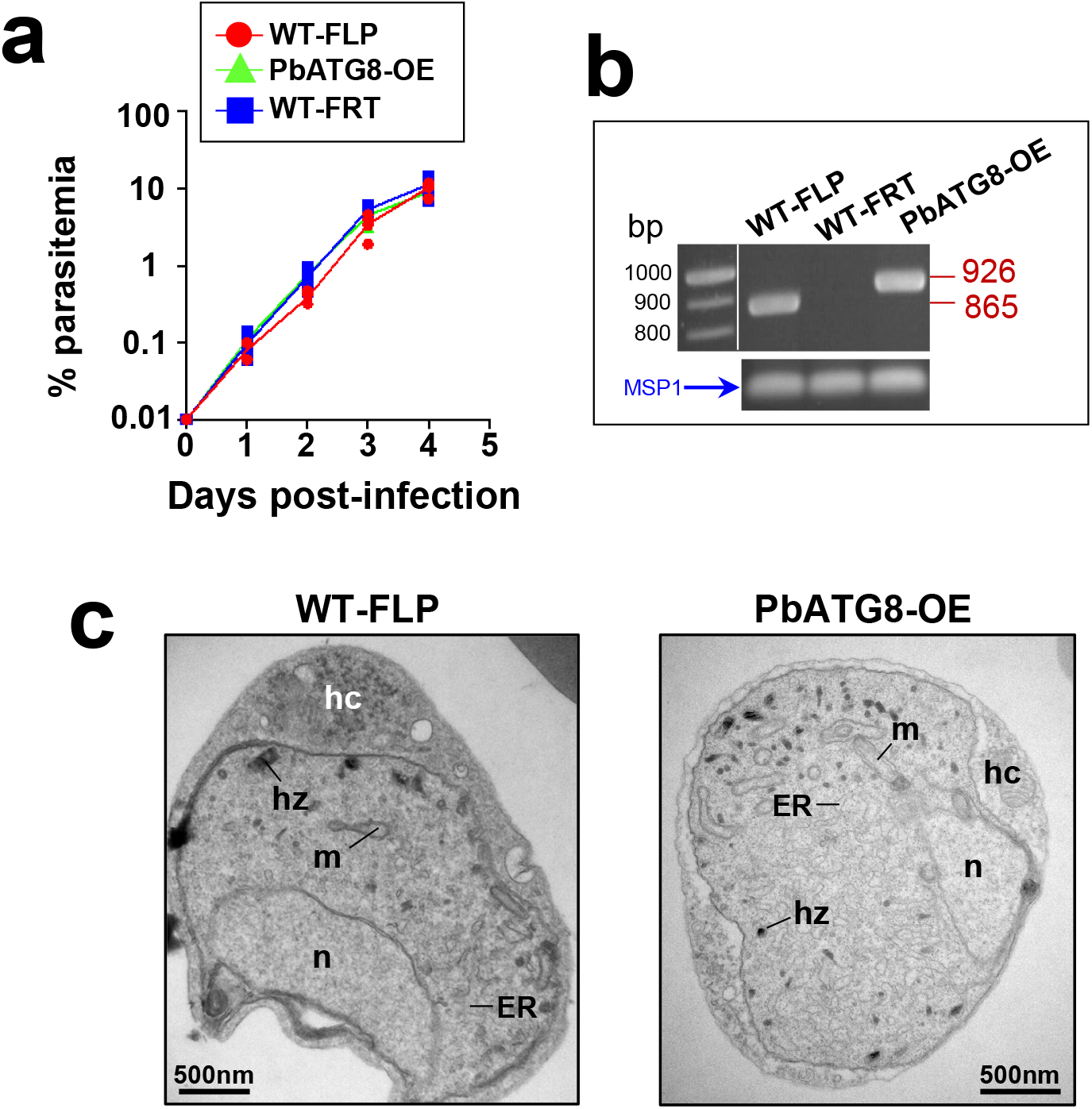
Genetic modification of the PbATG8-OE strain does not affect blood stage development. **a.** Naïve Swiss-Webster mice were i.v. infected with iRBC (0.01% final parasitemia) collected from donor mice infected with WT-FLP, WT-FRT (WT-FLP harboring the recombination construct but prior to recombination) or PbATG8-OE parasites. Parasitemia was monitored by bloodsmears every 24h for 4 days. (n=5 for each group; n.s. calculated by multiple t-test). **b.** Confirmation of the identity of each strain verified by PCR using primers encompassing part of ATG8-coding sequence (Forward-primer) and 3’UTR (Reverse-primer). This primer pair detects bands at 865-bp in WT-FLP and 926-bp in PbATG8-OE, and no amplification in WT-FRT parasites because of the disruption of the Reverse-primer due to the presence of the hDHFR cassette. *Plasmodium* MSP1 was used as positive control for the presence of parasite. **c.** Ultrastructure of PbATG8-OE parasites compared to WT-FLP. Representative electron micrographs of infected red blood cells with PbATG8-OE parasites or WT-FLP from 24 to 28 sections on different parasites. Mice were infected with WT-FLP or PbATG8-OE sporozoites, and blood from infected mice was collected 5- and 18-day post-inoculation. hc, host cell; hz, hemozoin; m, mitochondrion; n, nucleus.

### Immunization with PbATG8-OE-CVac protects mice in a dose-dependent manner

PbATG8-OE parasites remain in the liver longer than WT-FLP parasites due to their slow and defective liver stage development in hepatocytes. This finding suggests that the mutant exposes liver and blood stage antigens to the immune system for an extended period of time, making the liver infected with the mutant an immunologically favorable environment for the generation of anti-parasitic immunity in the host. Additionally, the mutation introduced in the 3’UTR of *Atg8* would presumably impose a selection pressure on the parasite, rendering it more vulnerable to another stress (e.g., drug exposure), which would impair its ability to develop resistance as compared to non-mutated parasites. We next wanted to investigate if PbATG8-OE parasites would likely be more sensitive to a chemoprophylaxis treatment, and hence an ideal antigenic constituent of a Chemoprophylaxis Vaccination regimen (CVac). To test if PbATG8-OE infection combined with drug treatment confers better immunity, we designed an immunization strategy using a stringent outbred mouse for vaccine studies as these animals represent the complex heterogeneity of human immunity and because they are difficult to protect with multiple RAS immunization^27^. We first determined the optimal number of sporozoite doses injected into mice that would be optimal to elicit a sterile protection as detailed in our experimental design (Fig. 3a). One, two or three doses of PbATG8-OE sporozoites with 10,000 parasites per dose were injected into Swiss-Webster mice along with chloroquine treatment. One week after the last immunization at D14 (21 days after the first dose), mice were challenged with 10,000 WT sporozoites. All mice that received a single immunization dose became positive by D5 while mice that were immunized with 2 or 3 doses of sporozoites, were protected at 80% and 100%, respectively.

**Figure 3.**
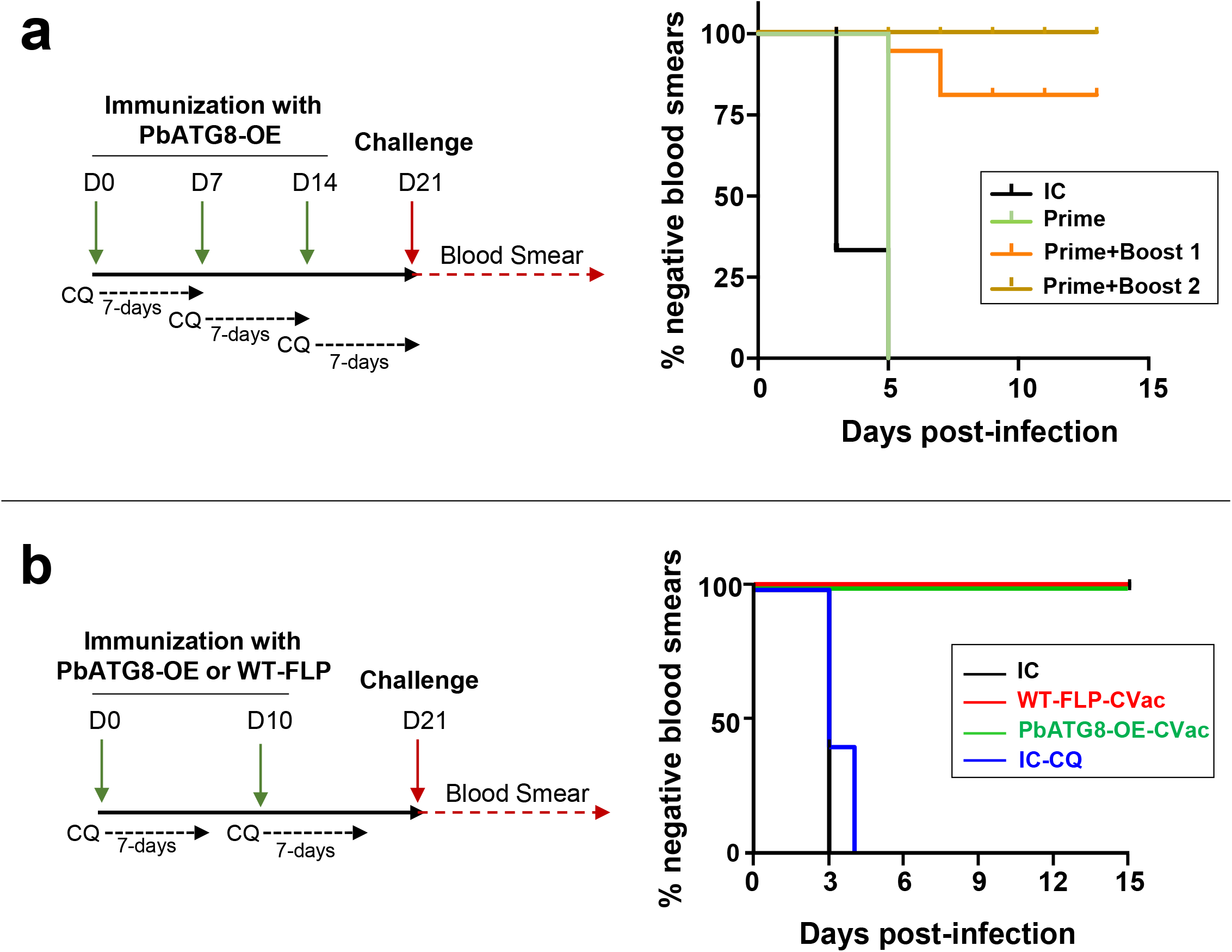
Immunization with PbATG8-OE-CVac parasites protects mice. **a.** Dose-dependent protection elicited by PbATG8-OE-CVac. Protocol for an immunization with PbATG8-OE parasites and challenge under CVac (PbATG8-OE-CVac). Swiss-Webster mice immunized with 10^4^ sporozoites from the PbATG8-OE strain and treated with chloroquine (0.8 mg/mouse) intraperitoneally for 7 days, starting from the day of sporozoite injection. Three separate groups (n=5) were immunized: first group received one dose of PbATG8-OE parasites at D0 [Prime]; second group received two doses 7 days apart [Prime + Boost 1]; third group received 3 doses 7 days apart [Prime + Boost 2]. All mice were challenged on D21 and parasitemia was monitored daily by thin blood smears, starting D3 post-challenge until D15. Kaplan-Meier survival plot showing days to blood stage infection after challenge with 10^4^ WT-Pb-ANKA sporozoites on D21. IC: Naïve infection control. (*p*=0.0039 for [IC] vs. [Prime]; *p*<0.0001 for [Prime vs. [Prime+Boost1]; *p*=0.0391 for [Prime+Boost1] vs. [Prime+Boost2], measured by log-rank Mantel Cox test. **b.** Time to blood stage infection during effector challenge. Protocol for immunization with PbATG8-OE or WT-FLP parasites and challenge under CVac. Swiss-Webster mice immunized with two doses of 2×10^4^ sporozoites from the WT-FLP or PbATG8-OE strain under 7 days of chloroquine treatment (0.8 mg/mouse) 10 days apart prior to challenge on D21. Parasitemia was monitored for 15 days starting from D3 post-challenge. Kaplan-Meier survival plot showing days to blood stage infection after challenge with 10^4^ WT-Pb-ANKA sporozoites. Two control groups included one infection control (IC) and one chloroquine control (IC-CQ) in which mice received chloroquine but no sporozoites. n.s. for [IC] vs. [CQ-IC]; *p*<0.01 for [CQ-IC] vs. [WT-FLP-CVac] and for [IC-CQ] vs. [PbATG8-CVac], measured by log-rank Mantel Cox test.

### PbATG8-OE-CVac parasites induce better primary and secondary memory response than WT-FLP-CVac parasites

Based on this finding in Fig. 3a, we devised a new immunization protocol in which two doses of sporozoites were injected into mice, but with a higher number of parasites (20,000) per dose administrated 10 days apart (Fig. 3b). We next sought to investigate to which extent PbATG8-OE parasites, due to their longer residency in the liver, would provide better protection than the parent parasite line in mice challenged with WT-FLP parasites. Following a challenge at D21, all mice immunized with either PbATG8-OE or WT-FLP parasites, were blood smear-negative for the entire period of observation (15 days post-challenge). In comparison, in the two control groups that include non-immunized mice, either treated with chloroquine (0.8 mg/mouse) or PBS control, all animals were infected at days 4 and 3, respectively. For challenges at D60 (primary memory response) or D80 (secondary memory response), a significantly better protection of mice was achieved for the mutant with 100% of blood smear-negative as compared to 60% for WT-FLP-CVac parasites, for both memory responses (Fig. 4). For long-term memory response probed with a challenge at D180, PbATG8-OE-CVac parasites conferred 50% protection as compared to 30% for WT-FLP-CVac parasites. However, the difference was not statistically significant between the two strains though observed in two independent experiments (*p*=0.233 and *p*=0.63).

**Figure 4.**
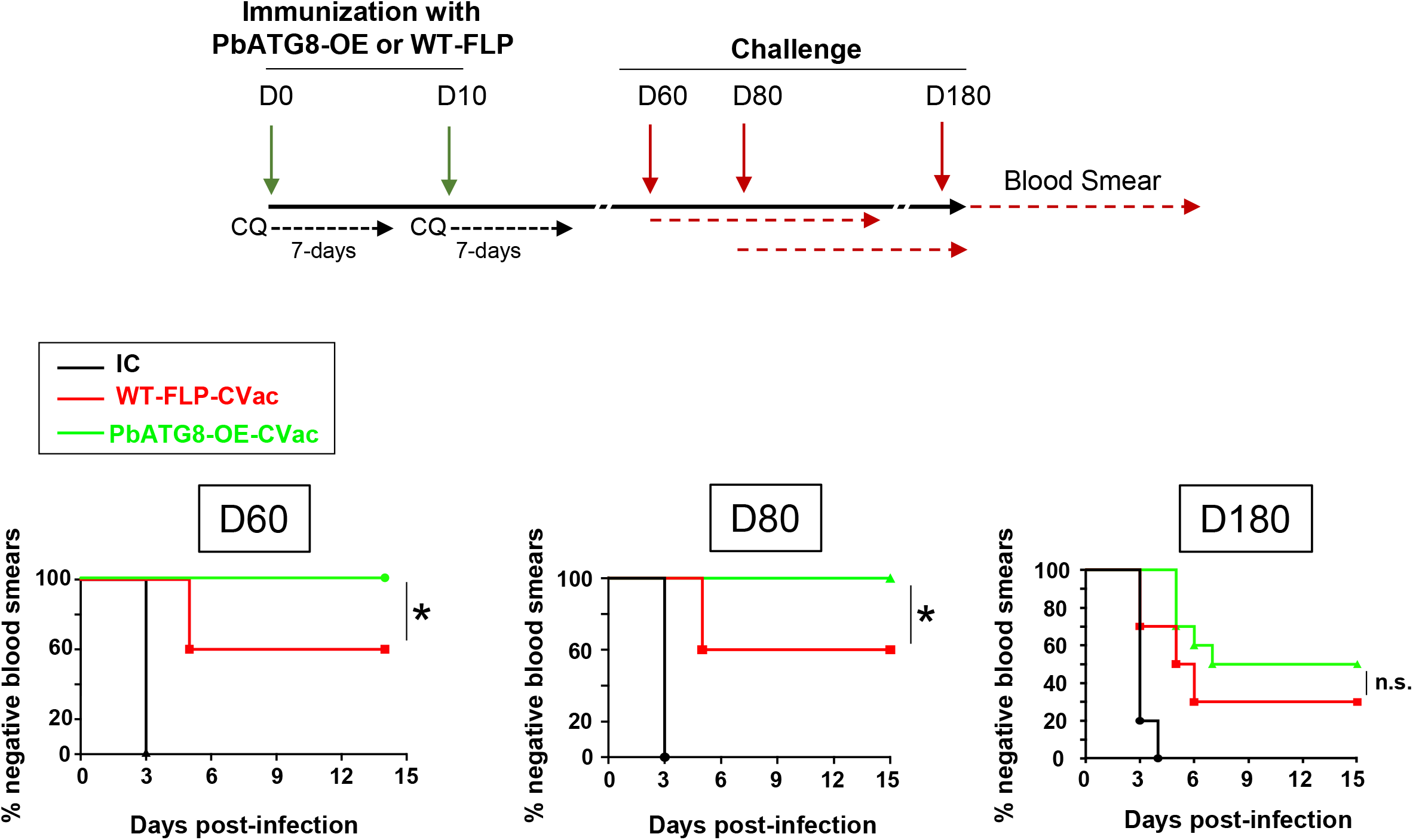
PbATG8-OE-CVac parasites elicits long-term memory protection in mice. Protocol for immunization with PbATG8-OE or WT-FLP parasites for memory challenge. Swiss-Webster mice immunized with two doses of 2×10^4^ sporozoites of WT-FLP or PbATG8-OE parasites 10 days apart. Immunized mice were treated with chloroquine (0.8 mg/mouse) for 7 days starting with sporozoite immunization. Mice were then divided into three groups and challenged with 10^4^ WT-Pb-ANKA sporozoites at memory time points D60 and D80 and D180 post-immunization prior to parasitemia determination by Giemsa stained thin blood smears from tail snip from D3 post-challenge until D15. Kaplan-Meier survival plots showing days to blood stage infection after challenge on D60 (a), D80 (b) and six months (c). Data are representative of experiment done in duplicate with 5 mice in each group for D60 and D80, and 10 mice in each group for six-month protection analysis. IC: non-immunized infection control (n=5 each time). *, *p*<0.05, measured by log-rank Mantel Cox test.

We further verified the sterile protection in PbATG8-OE-CVac-immunized mice upon a challenge at D80. Compared to WT-FLP-CVac-immunized mice that came blood stage-positive at day 5, and increased over time, the parasitemia remained null in PbATG8-OE-CVac-immunized mice (Fig. 5a). To further confirm the total protection of PbATG8-OE-CVac-immunized mice observed at D80 post-immunization, 100 μl of blood from each mouse immunized with either mutant or parental sporozoites and challenged, were transferred on D15 post-challenge to naïve receiver mice. Blood stage infection in mice was then monitored for 2 weeks. All mice that received blood from PbATG8-OE-CVac-immunized mice were blood smear-negative compared to 60% that received blood from WT-FLP-CVac-immunized mice (Fig. 5b). Mouse blood infection was further verified by amplification of ~120 bp of Pb-18S rRNA gene (Fig. 5c), and data were in accordance to results in Fig. 5b. A separate group of mice immunized with PbATG8-OE or WT-FLP parasites were also challenged by mosquito bite in which WT-FLP-CVac-immunized mice received 3 to 5 infectious bites and PbATG8-OE-CVac-immunized mice received 6 to 8. Mice from the two groups were completely protected from the mosquito bite challenge, whereas all naïve mice became blood stage-positive from days 5 to 10 post-challenge (Fig. 5d).

**Figure 5.**
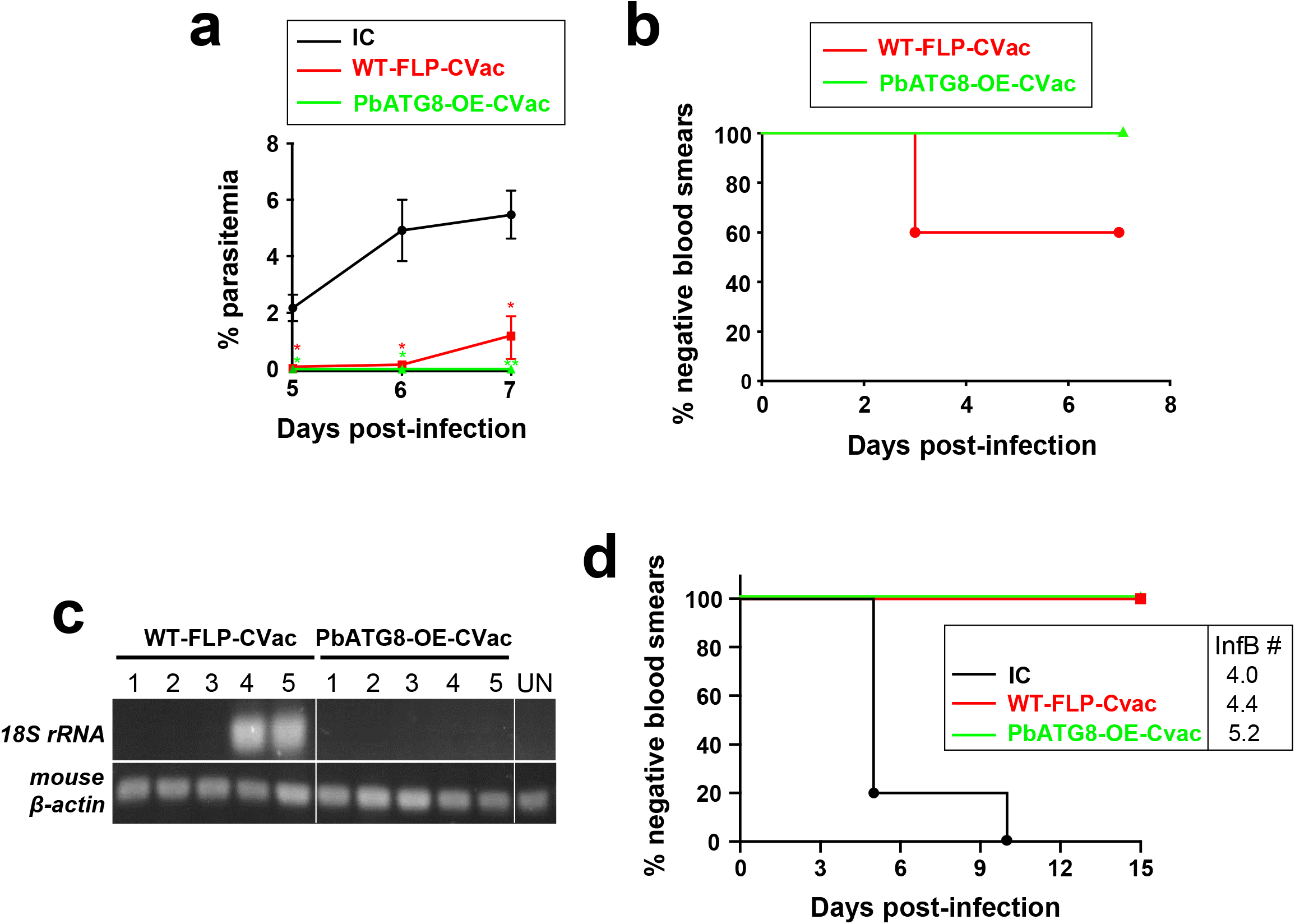
PbATG8-OE-CVac-immunized mice are sterilely protected at memory time points. **a.** Percent parasitemia during memory challenge. Swiss-Webster mice immunized with 2×10^4^ WT-FLP-CVac or PbATG8-OE-CVac sporozoites were challenged on D80 post-immunization with 10^4^ WT-Pb-ANKA sporozoites to monitor parasitemia from D5 post-challenge. Data are means ± SD of 5 mice in each group (*, *p*<0.005; **, *p*<0.0001, measured by multiple t-test). **b.** Prepatent period during memory challenge. Blood (0.1 ml) from challenged (on D80) mice were transferred on D15 post-challenge to recipient naïve mice to monitor the parasitemia for 15 days (*p*<0.05, measured by log-rank Mantel Cox test). **c.** PCR verification on blood from the infected mice at D80, with the amplification of 130 bp of Pb-18S rRNA gene and mouse β-actin (control). Genomic DNA isolated from uninfected mice (UN) was used as a negative control. **d.** Challenge with mosquito bites. A separate group of immunized mice (n=5 per group) were challenged WT-Pb-ANKA sporozoites through mosquito bite at D80. Mice from IC group received an average of 4 infectious bites (InfB), WT-FLP-CVac-immunized mice 4.4 InfB and PbATG8-OE-CVac-immunized mice 5.2 InfB.

### Protection conferred by PbATG8-OE-CVac parasites is predominantly CD8^+^T cell-dependent and specific to pre-erythrocytic stages

We next examined the potential protective role of CD8^+^ and CD4^+^T cells in mice immunized with PbATG8-OE parasites by selective antibody-mediated depletion after immunization of outbred Swiss-Webster mice. It is known that CD8^+^ T cells contribute significantly towards anti-malarial immunity, in clearing *Plasmodium*-infected hepatocytes, in a process that is IFNγ-dependent^32–34^. Expectedly, CD8^+^T cell immunodepletion results in 80% loss of protection in mice immunized with PbATG8-OE parasites, in contrast to 20% loss of protection under condition of CD4^+^T cell immunodepletion (Fig. 6).

**Figure 6.**
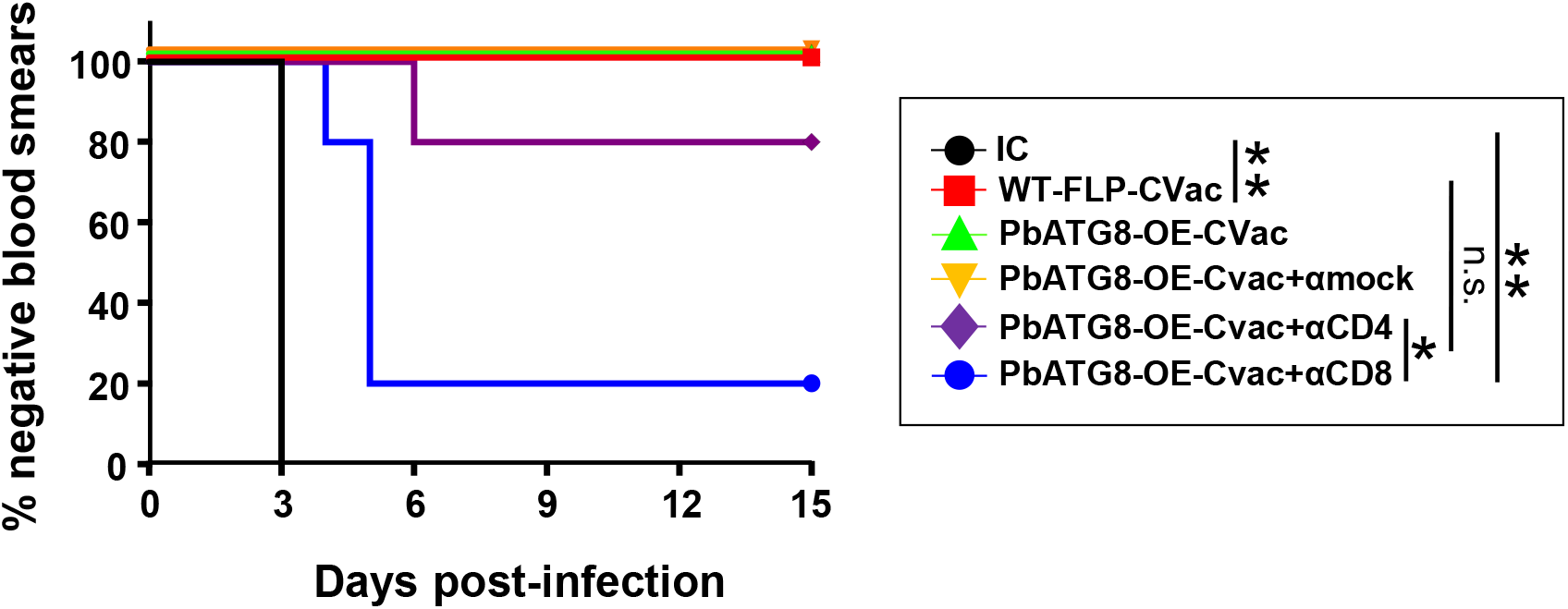
CD8^+^T-cell depletion post-immunization reduces protective response elicited by PbATG8-OE-CVac parasites. Kaplan-Meier survival plot showing days to blood stage infection. Swiss-Webster mice were immunized with 2×20,000 WT-FLP-CVac or PbATG8-OE-CVac sporozoites. PbATG8-OE-CVac-immunized mice were divided into 4 groups (n=5 in each group). Three groups of mice received intraperitoneally 500 μg of antibodies per dose: αCD8 (clone 2.43), αCD4 (clone GK1.5) or mock isotype control IgG (LTF-2). Antibody treatment were done twice, 24h prior to challenge and on the day of challenge. Parasitemia was monitored daily following the challenge with 10^4^ WT Pb-ANKA sporozoites, *, *p*<0.05; **, *p*<0.01, measured by log-rank Mantel Cox test. Data are representative of one experiment done twice.

In order to verify if the protective immunity is specific to the pre-erythrocytic stage or is extended to the blood stage, we devised two immunization protocols to assess the effector and memory response in PbATG8-OE-CVac-immunized mice following challenge with iRBC at D21 (effector response) and D80 (memory response). The protective immunity rendered by PbATG8-OE-CVac and WT-FLP-CVac parasites did not protect against any of the two blood stage challenges, even with a high sporozoite antigen dose used for the effector response (Fig. S1a-b). These data suggest that the CD8^+^T cell-dependent protection mediated by PbATG8-OE-CVac parasites is likely to be specific to pre-erythrocytic stages.

### Protection rendered by PbATG8-OE-CVac parasites is largely independent of IFN-γ-mediated immune response

It has been well-documented that a CD8^+^T cell-mediated protection in immunized mice with sporozoites (e.g., RAS or GAP) is largely IFN-γ dependent^5,9,34,35.^ To examine the contribution of IFN-γ, we immunized C57BL/6J WT mice and C57BL/6J KO mice for either IFN-γ or IFN-γ receptor with PbATG8-OE-CVac parasites prior to challenge with WT sporozoites expressing luciferase. Liver stage burden was measured by in vivo bioluminescence imaging 42 h after sporozoite challenge. All non-immunized mice used as controls exhibited a strong bioluminescence signal in their liver (Fig. 7a). Interestingly, upon immunization of IFN-γ KO and IFN-γ receptor KO mice with PbATG8-OE-CVac and subsequent challenge with WT parasites, the bioluminescence signals were close to background levels in all immunized IFN-γ receptor KO mice and 80% of immunized IFN-γ KO mice. Based on quantification of bioluminescence signal intensity, no significant difference in the parasite load in the liver of the KO mice was observed with PbATG8-OE-CVac-immunized WT mice (Fig. 7b). By monitoring the subsequent parasitemia, 20% of immunized IFN-γ KO mice became blood stage-positive after 10 days. However, no statistically significant difference was calculated between IFN-γ receptor KO and WT-immunized mice (Fig. 7c). Nevertheless, liver parasite burden and parasitemia were significantly less than observed with non-immunized mice. No loss of protection in the KO mice suggests CD8^+^ T cells may be rendering its protective action through effector/s other than IFN-γ.

**Figure 7.**
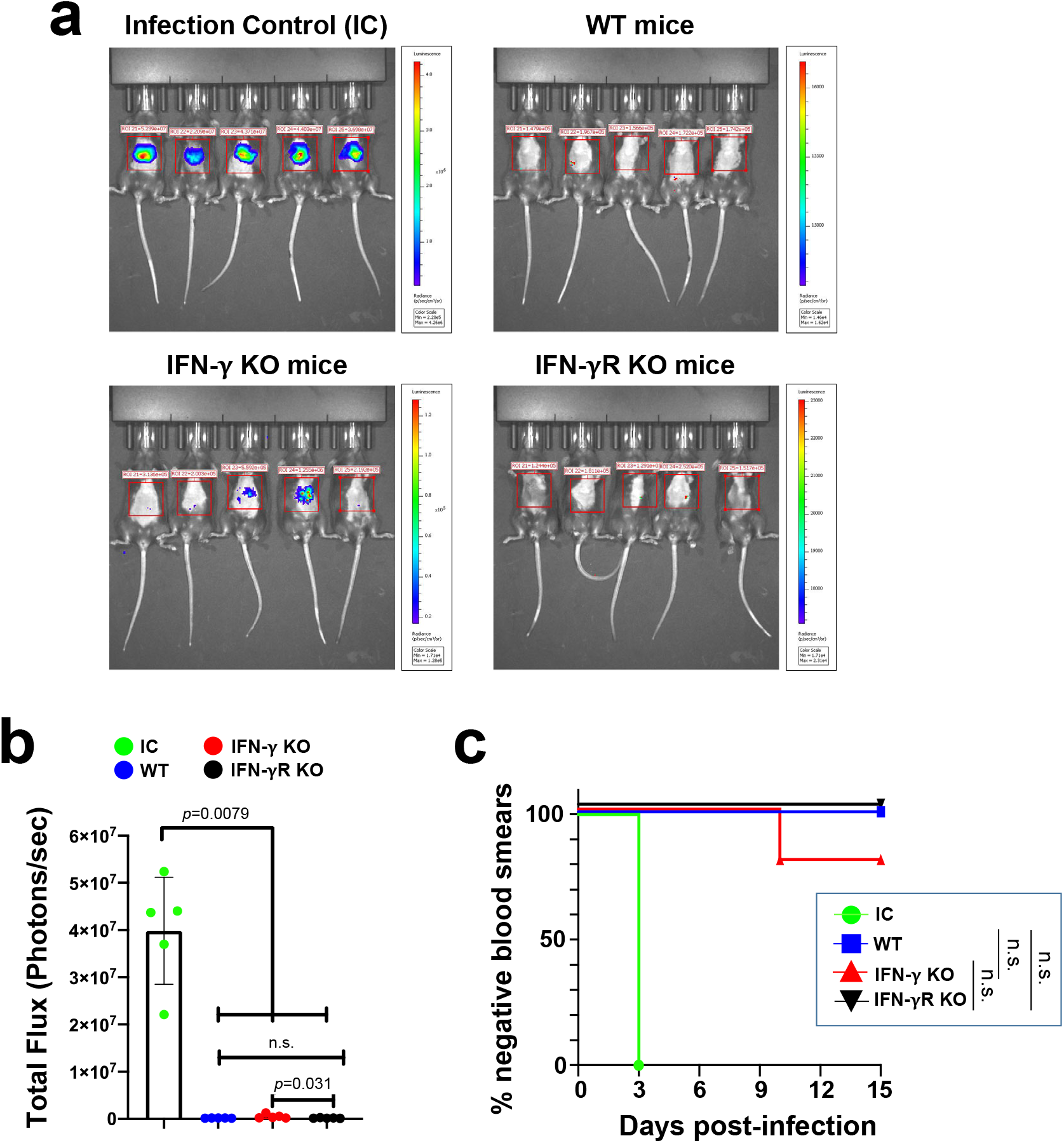
IFN-γ is not required for protection rendered by PbATG8-OE-CVac parasites. **a-b.** Bioluminescence imaging of mice infected with Pb-GFP-Luc parasites to assess parasite burden in the liver and quantification. Absolute luminescence was measured as photon/second in infection control mice and PbATG8-OE-CVac-immunized animals: WT mice, IFN-γ-KO mice or IFN-γ R-KO mice. Data representative of one experiment done twice with 5 mice in each group, are means ± SD (values in photons/sec were 169,520 ± 18,723 for WT mice; 509,460 ± 440,595 for IFN-γ-KO mice; 167,660 ± 52,218 for IFN-γ R-KO mice). Significance was determined by Mann-Whitney test. **c.** Kaplan-Meier survival plot showing days to blood stage infection. WT: C57Bl/6J mice immunized with PbATG8-OE-CVac parasites. n.s. *p*>0.05, measured by log-rank Mantel Cox test.

### PbATG8-OE-CVac parasites trigger a broad but PbCSP-specific anti-sporozoite antibody response

We also assessed potential humoral responses in the protection of mice immunized with PbATG8-OE-CVac compared to WT-FLP-CVac-immunized mice. Sera from naïve or immunized mice as described in Fig. 3b were collected 4 weeks after the last immunization, pooled from different mice and diluted for IFA on iRBC, extracellular sporozoites and liver forms in Hepa1-6 at different time points of infection (24 h, 44 h and 67 h) representing different developmental stages during liver stage infection. IFA using sera for immunized mice illustrate a strong fluorescence staining on all parasitic stages, without any discernable difference between parasite labeling after exposure to sera from mice immunized either with mutant or parental parasites (Fig. 8a). These observations indicate that immunized mice from either group have produced antibodies that recognize antigens commonly expressed in all *Plasmodium* stages. Evaluation of a functional antibody response was performed using a Sporozoite Neutralization Assay (SNA), in which freshly dissected salivary gland sporozoites were incubated ex vivo with the same sera collected from immune or naïve mice that were used for IFA. Serum-treated sporozoites were then injected intradermally into naïve mice (to mimic a natural route of sporozoite infection by mosquito bite), to monitor the development of blood stage infection by blood smears. All mice infected with sporozoites pre-exposed to antisera from PbATG8-OE-CVac- or WT-FLP-CVac-immunized mice were refractory to blood stage infection (Fig. 8b). This finding reveals that the antibodies produced by the immunized mice prevent infection, and thus efficiently protect.

**Figure 8.**
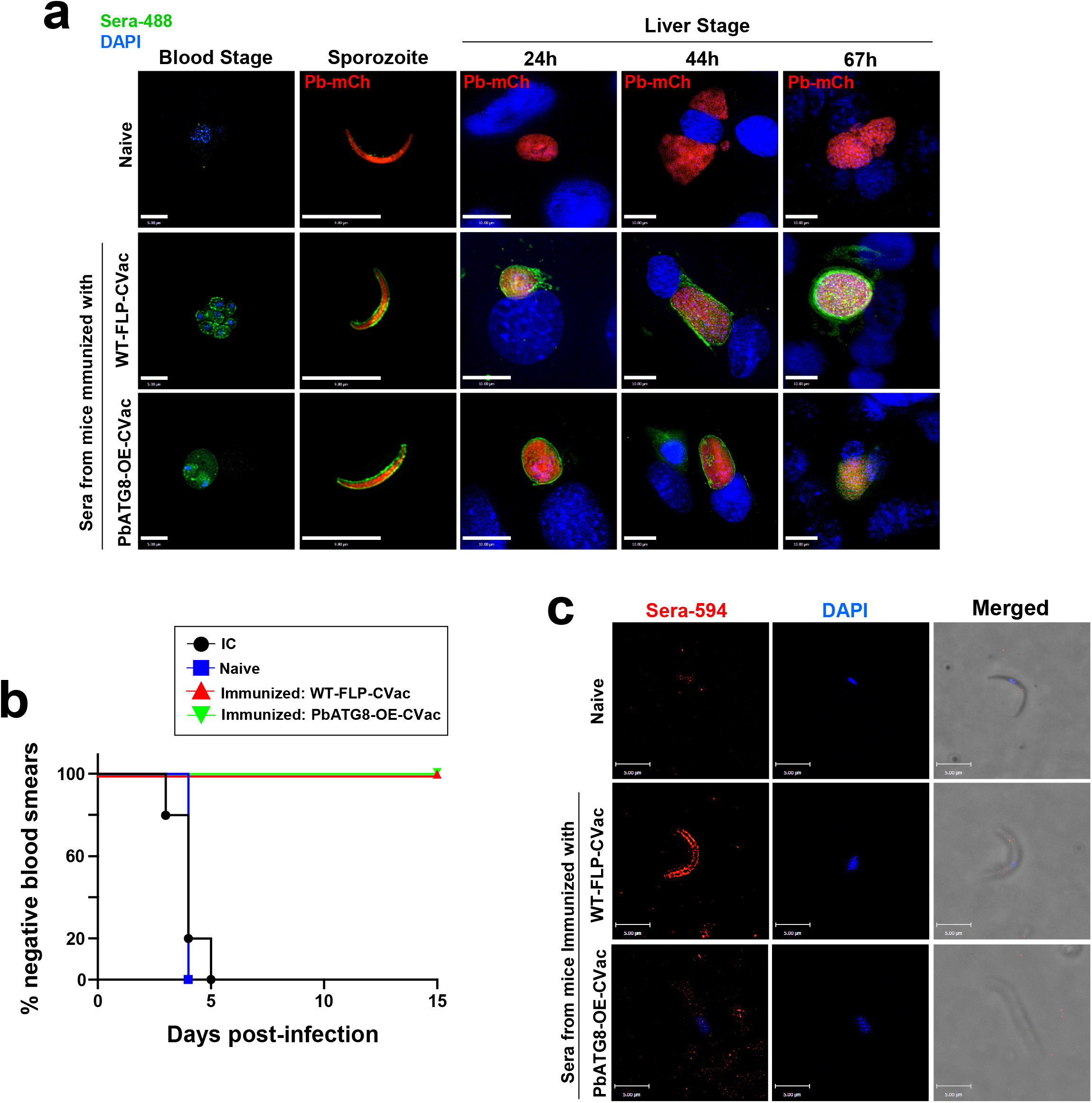
Antibody response generated by PbATG8-OE-CVac parasites is protective. **a.** IFA using diluted sera from naïve, WT-FLP-CVac or PbATG8-OE-CVac mice performed on mixed blood forms, *P. berghei* expressing mCherry (Pb-mCh) sporozoites or liver forms at different time points. Mice were immunized as described in Fig. 3b and sera used for IFA were collected 4 weeks after the last immunization. Fluorescence signal was detected by Alexa-488 conjugated secondary anti-mouse-IgG. Nucleus was stained with DAPI. Scale bars, 10 μm. **b.** Sporozoite neutralization assay. 2×10^4^ WT-Pb-ANKA sporozoites were incubated on ice for 45 min with 1:6 diluted sera from naïve or immunized mice, and the sporozoite-antibody mixture was injected intradermally to naïve Swiss-Webster mice and parasitemia was monitored for 15 days. **c.** IFA using diluted sera from naïve or immunized mice on non-permeabilized air-dried PfCSP-expressing *P. berghei* sporozoites. Alexa-594 conjugated secondary anti-mouse IgG were used to detect antibody signal at the sporozoite surface. Scale bars, 5 μm.

It is well-described that anti-sporozoite humoral immunity generated by whole sporozoite vaccine is mostly directed towards CSP^36–38^. We next wanted to explore the targets of the anti-sporozoite antibody associated with sterile protective immunity in mice immunized with PbATG8-OE-CVac. IFA were performed using sera collected from PbATG8-OE-CVac- or WT-FLP-CVac-immunized mice on non-permeabilized transgenic *P. berghei* sporozoites expressing *P. falciparum*-CSP^22^, in place of the *P. berghei*-CSP. In contrast to sporozoites in Fig. 8a, no fluorescence signal was detected at the surface of PfCSP-*P. berghei* sporozoites incubated with antiserum from PbATG8-OE-CVac-immunized mice (Fig. 8c). This observation suggests that this antiserum contains mostly anti-PbCSP antibodies, and no or very few antibodies against sporozoite surface antigens other than PbCSP, resulting in no detectable signal by IFA. In comparison, the surface of PfCSP-*P. berghei* sporozoites exposed to antiserum from mice immunized with parental parasites was uniformly labeled, indicative of the presence of anti-sporozoite antibody target/s other than PbCSP. These data reveal that immunization with PbATG8-OE-CVac may engender mainly a PbCSP-specific antibody response, which may confer protection.

### PbATG8-OE-CVac parasites induce a better memory T-cell response than WT-FLP-CVac parasites

We show that PbATG8-OE-CVac parasites confer better memory response and long-term protection than WT-FLP-CVac parasites (Figs. 4 and 5). To examine the qualitative difference among the antigen-exposed CD8^+^ T cells between PbATG8-OE-CVac- and WT-FLP-CVac-immunized mice, we investigated the anti-parasite specific CD8^+^ T cells population in an approach using the CD8α^lo^CD11a^hi^ surrogate activation marker^7,27^. T cells expressing low levels of CD8α and high levels of CD11a represent antigen-experienced cells. The sensitivity and amplitude of the CD8α^lo^CD11a^hi^ T cells to mount antigen response was monitored by FACS analysis of peripheral blood collected from naïve mice and mice immunized with PbATG8-OE-CVac or WT-FLP-CVac. The number of circulating primed CD8α^lo^CD11a^hi^ T cells was almost double in PbATG8-OE-CVac-immunized mice than in mice immunized with parental parasites (Fig. 9a). Three-month post-immunization, the decline in this T cell population from the level that was observed after final immunization was less pronounced in the blood of mice immunized with PbATG8-CVac parasites compared to WT-FLP-CVac-immunized mice (Fig. 9b). These data suggest that CD8α^lo^CD11a^hi^ T cells are more abundant and/or subsist for longer periods of time in PbATG8-OE-CVac-immunized mice, reflecting their superior efficiency to control subsequent infection during challenge.

**Figure 9.**
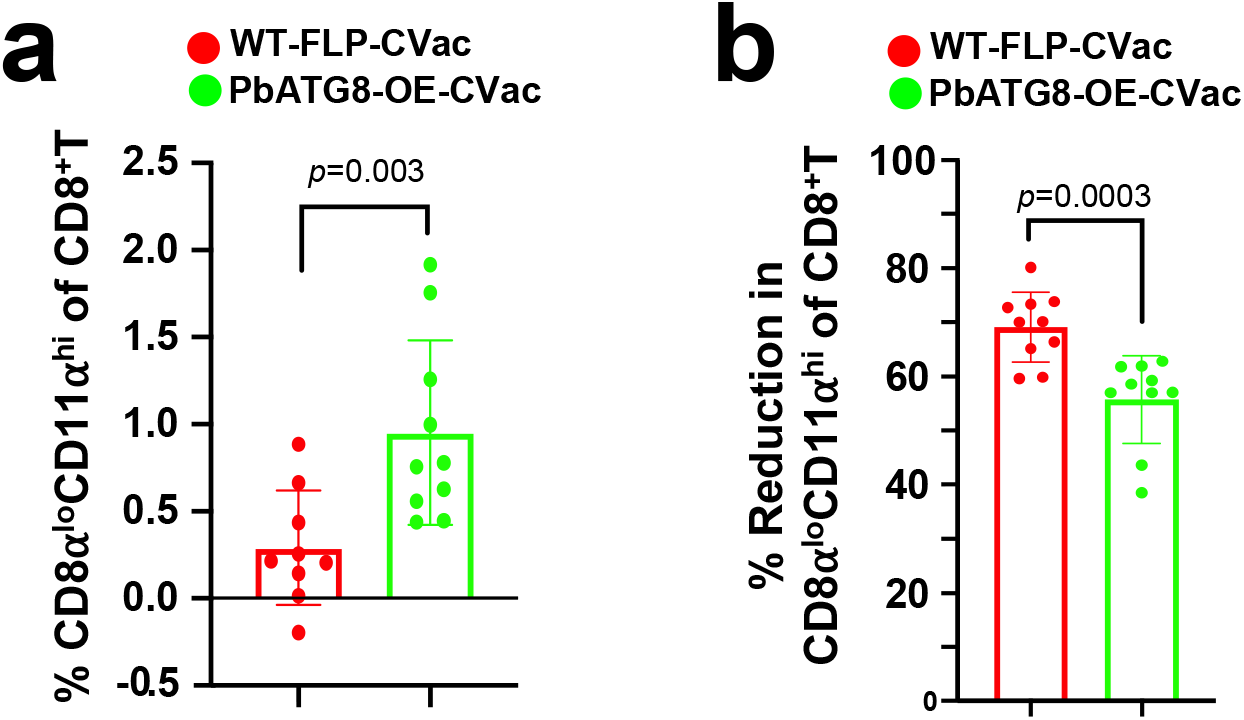
Antigen-exposed CD8^+^T cells have unique dynamics upon PbATG8-OE-CVac immunization. **a.** FACS analysis for CD8α^lo^CD11a^hi^-exposed CD8^+^T cells in peripheral blood. Blood of C57BL/6J mice was analyzed 5 days prior to (preimmune) and 7 days after immunization with 2×10^4^ WT-FLP-CVac or PbATG8-OE-CVac sporozoites for the presence of CD8α^lo^CD11a^hi^ T cells by FACS. Data expressed as % of total CD8^+^T cells, are means ± SD (n=10 mice in each group). Statistical significance was determined by Mann-Whitney test. **b.** The graph represents the contraction (% reduction) in the CD8α^lo^CD11a^hi^ population in peripheral blood of immunized mice 3 months after last immunization, normalized to post-boost population of the CD8α^lo^CD11a^hi^ T cell population. Means ± SD (10 mice in each group). Statistical significance was determined by Mann-Whitney test.

## Discussion

Vaccination using WT sporozoites in conjunction with an anti-malarial drug administered as prophylaxis (CVac) has shown enhanced efficacy as a whole organism vaccine because the parasite progresses through all liver developmental stages, a potent immune response is triggered against a significant biomass of immunogens and the risk of acute blood stage infection is eliminated with drug treatment. However, a live sporozoite-CVac vaccine protocol remains difficult to control in the human population as it crucially relies on the drug regimen and proper use by individuals, and could lead to sporadic development of malaria symptoms in case of failure. As a safer alternative, replacement of fully infectious sporozoites by a genetically attenuated *Plasmodium* strain that can invade but poorly replicate within hepatocytes may constitute a promising direction for whole sporozoite vaccine. In this study, we reported that a liver stage-specific conditional mutant *Plasmodium berghei* (PbATG8-OE) when used as a whole-sporozoite vaccine under CVac regiment with chloroquine, provides better long-term protection than parental parasites under the same immunization protocol. Immunization of mice with PbATG8-OE-CVac parasites results in a significantly higher population of the antigen-experienced CD8^+^T cells during priming that remains comparatively stable, even 3 months after immunization. Interestingly, PbATG8-OE-CVac parasites generate a protective CD8^+^ T cell-mediated cellular response against liver-stage that is minimally dependent on IFN-γ, unlike many other whole sporozoite vaccines that trigger IFN-γ for protection, such as RAS^35^, WT-CVac with chloroquine or trimethoprim-sulfamethoxazole^9,25,35,36^ and GAP^40^.

The persistence of a parasite in the liver due to a replication deficiency in hepatocytes would induce a very strong immune response as previous observations suggest that persistence of liver stage antigens is important for optimal CD8^+^ T cell response against *Plasmodium* liver forms^41,42^. In addition to the large repertoire of antigens from late GAP presented to T cells, expression of hepatic merozoite antigens may trigger cross-stage protection against blood forms. Late liver stage-arresting replication-competent strains engineered so far are based on targeted gene deletion. As an alternative, we generated a liver stage-specific conditional mutant that overexpresses the essential gene *Atg8*, resulting in dysregulated autophagy machinery of the ATG8 ubiquitination system^18^. The stable insertion of a CG-rich nucleotide sequence in the 3’ UTR of the *Atg8* gene (to generate more stable secondary RNA structures^43,44^) successfully leads to higher levels of Atg8 transcripts, up to a ~3-fold increase at 68 h post-infection, and is associated with doubling the expression of PbATG8 at the protein level. Like observed for LC3 in mammalian cells^45^, interfering with ATG8 function and regulation through altered expression levels is detrimental for *Plasmodium* liver forms. Indeed, PbATG8-OE parasites suffer from severe differentiation defects and replication delays late during schizogony in hepatocytes both in vitro and in vivo^18^ (this study). Defective differentiation events in PbATG8-OE parasites include impaired elimination of obsolete sporozoite organelles (e.g., micronemes) by exophagy. ATG8 is expressed on the endosymbiotic apicoplastic organelle and ATG8-containing membranes of the apicoplast serve as lipid providers for the formation of autophagosomal structures that enclose micronemes prior to degradation^18,19^. In PbATG8-OE parasites, the apicoplast shows several morphological defects, which likely does impact liver form survival. Importantly, *Plasmodium falciparum* also expresses ATG8 on the apicoplast during liver residency, and PfATG8 displays 88% identity with PbATG8, making the exploitation of PfATG8-OE parasites as whole-parasite vaccines relevant for human malaria.

During a natural infection with *Plasmodium*-infected mosquitoes, liver stage malaria affords only a short window of opportunity to mount protective CD8^+^T cell responses, as only a few dozen to a hundred sporozoites are inoculated into the host dermis, leading ultimately to a very low number of infected hepatocytes before release of merozoites into the bloodstream (about 7 days in humans and 2 days in rodents)^33^. This limitation can be overcome if sufficient antigen-specific CD8^+^ T cell responses are triggered against liver stages, like through the needle-delivery of liver stage-arresting GAP at high sporozoite doses (i.e., in the range of thousands of sporozoites, equivalent to a thousand mosquito bites). Compared to parental parasites, PbATG8-OE parasites stay significantly longer in the liver, up to 12 days. After a prolonged period of development, most PbATG8-OE EEF are cleared from hepatocytes before reaching the merosomal stage. Among the EEF that survive, some could manage to produce more or less infectious merozoites, leading to null^18^ or low-frequency breakthrough infections (this study). Overall, the mutant may be exploited as a potentially interesting whole-organism vaccine under CVac regimen.

Protective immunity targeted against intrahepatic parasites is complex and multifactorial. Many vaccines that exploit T cell-mediated immunity to malaria establish the importance of T cells directed against liver stage antigens and antibodies mounted against sporozoite surface proteins. Studies based on experimental malaria models (rodents, non-human primates) or on humans reported that CD8^+^ T cells are important for the elimination of *Plasmodium*-infected hepatocytes, with IFN-γ being the critical effector molecule^27,46–50^. IFN-γ has multiple effects on innate and adaptive immune responses, such as proinflammatory priming of Toll-like receptor responses and upregulation of MHC class I and II expression^51^. If CD8^+^ T cells are the primary effector cells in protective immunity, CD4^+^T cells that recognize parasite-derived/class II MHC molecule complexes on the surface of infected hepatocytes also participate in the effector mechanisms against malaria liver stages^52^. Additionally, the subsets of CD4^+^T cells that secrete IL-4 or IL-12 provide help in inducing the CD8^+^ T cell responses and for optimal CD8^+^T cell effector activities, respectively, suggesting a CD4-CD8 cross-talk for the development of anti-malaria protective immunity^53^. Thus, the main core elements against infected hepatocytes consist of CD8^+^ and/or CD4^+^T cells and IFN-γ-mediated responses. In our system using PbAGT8-OE-CVac, we showed in conformity with our previous reports, that CD8^+^T cells are important effectors for sterile protection and long-term memory response against infectious WT challenges. Interestingly, IFN-γ has a negligible contribution to immune protection in our model, contrasting with other studies on whole organism immunization. Indeed, IFN-γ receptor KO mice and 80% of IFN-γ KO mice immunized with PbATG8-OE parasites do not develop any blood infection after challenge, indicating protection even in the absence of IFN-γ activities. The slight difference in infectivity observed between mice lacking IFN-γ or IFN-γ receptor in our system may be due to variability in the immune and inflammatory status of the 2 KO mice. To this point, a study reports a more intense inflammation in IFN-γ receptor KO mice, with higher levels of proinflammatory cytokines and MHC proteins, in the context of sindbis virus infection^54^. The immune protection elicited by WT-CVac or GAP is mostly, if not entirely IFN-γ-dependent ^9,40^. In one study using a *Plasmodium yoelii* GAP (PyGAP) vaccine, sterile protection fully dependent on CD8^+^ T lymphocytes and partially on IFN-γ has been observed: 50% of mice (either IFN-γ KO or IFN-γ-immunodepleted mice) are protected from parasitemia^55^. In PyGAP-immunized mice, infected hepatocytes undergo apoptosis induced by CD8^+^T cells through pore formation by perforin. The mechanism of protection and effectors involved in the cytolytic activity of CD8^+^T cells triggered by PbATG8-OE parasites remain to be identified. In the absence of IFN-γ, such effectors (perforin/granzyme, or others) may serve as immunological correlates of protection (CoP) for anti-malaria vaccines.

Immune protection elicited by PbAGT8-OE-CVac parasites is mostly CD4^+^T cell-independent. From many studies on malaria rodent models, it has been established that CD4^+^T cells play an important role during the initial immune response against liver stage^56^, however, these cells are not actively involved in the protection during the effector immune response, more particularly during immunization with whole sporozoites^57–60^. For other protocols of immunization in which CD4^+^T cells have been identified as efficient contributors to the effector immune response, the protective role of CD4^+^T cells remains variable, depending on the type of the immunogen as well as the parasite-host combination^32,60,61^. Thus, like for many GAP vaccines, the protection induced by PbAGT8-OE-CVac parasites also depends critically on CD8^+^T cells, compared to CD4^+^T cells. This suggests a minimal contribution of CD4^+^T-cell-mediated for antibody production and CD4^+^ MHC class II-restricted T cells to cytolytic activitied against parasite-infected hepatocytes, and no CD4-CD8 cross-talk for the development of protective immunity, particularly during the effector phase of the immune response.

Anti-sporozoite antibody response is a strong contributor towards protective immunity^36,38,62^. In our Sporozoite Neutralization Assay, all mice were protected following the inoculation of sporozoites with sera from mice immunized with PbAGT8-OE-CVac parasites, indicative of a strong anti-sporozoite antibody response. This finding contrasts with another study using WT-CVac regimen with live *P. yoelii* sporozoite immunization under trimethoprim-sulfamethoxazole prophylaxis, in which no anti-sporozoite antibody response was observed^25^. However, in this CVac protocol with *P. yoelii*, antiserum-treated sporozoites were i.v. injection into the naïve mice and thus would have bypassed the locomotive requirement in the skin. Indeed, it has been established that the neutralizing capacity of circulating antibodies (e.g., by decreasing sporozoite motility and invasion) is greater at the inoculation site than in the bloodstream^63^.

Antibodies generated by mice immunized with PbAGT8-OE-CVac parasites bind to a plethora of antigens expressed by blood and liver forms, though the majority of their target antigens as well as the protective capacity of these antibodies are unknown. *P. berghei*-mCherry sporozoites are coated by protective antibodies from mice immunized with either PbAGT8-OE-CVac or WT-FLP parasites. However, the composition of the mouse antisera upon mutant or parental parasite immunization differs, with the production of antibodies mainly specific against PbCSP by mice immunized with PbAGT8-OE-CVac parasites. As the major surface protein of sporozoites, CSP is the target of sporozoite-induced CD8^+^T cell responses, and CD8^+^T cells against immunodominant CD8^+^T cell epitopes on CSP are able to eliminate infected hepatocytes (reviewed in ^64^). One possibility is that immunization with PbAGT8-OE parasites may confer protection through the activation and expansion of CD8^+^T cells that efficiently target PbCSP on sporozoites. On the contrary, antibodies from mice immunized with WT-FLP-CVac parasites may bind PbCSP but also recognize other protein/s at the surface of sporozoites, possibly SSP2/TRAP, HEP17 or Exp-1 that are known to be antigenic targets of CD8^+^T cell responses against *P. yoelii* or *P. falciparum* challenge as summarized in^64^.

PbATG8-OE-CVac parasites trigger a significantly higher CD8α^lo^CD11a^hi^ T cell population in immunized mice than parental-CVac parasites after one dose of immunization. This expansion of antigen-experienced T cells could be due to rapid expansion of early PbATG8-OE liver forms providing a greater exposure to parasite antigens, however, PbATG8-OE parasites develop normally at the onset of infection in hepatocytes. Alternatively, the mutant may be deficient in avoiding normal host defense mechanisms. Prior to multiplication in erythrocytes, parasites need to overcome many hepatic immune assaults and escape incognito from hepatocytes (reviewed in ^65^). The replication defects of PbATG8-OE-CVac liver forms leading to dead parasites may be associated with increased processing and presentation of parasite-derived peptides to MCH class I on the surface of infected hepatocytes for cytotoxic CD8^+^T cells activity. The usurpation of the plasma membrane of the host hepatocyte to form the merosomal membrane enwrapping hepatic merozoites is an efficient parasitic strategy to ensure the protection of merozoites from host phagocytes. To this point, the aberrant formation of PbATG8-OE merozoites may be associated with defects in the merosomal membrane, with the risk of parasite exposure to the immune system.

As opposed to previously reported whole sporozoite vaccines such as RAS, GAP and WT-CVac, PbATG8-OE-CVac parasite confers 100% protection at two memory time points (D60 and D80 post-immunization) in a clinically relevant outbred mouse model. The induction of long-term protection elicited by PbATG8-OE-CVac parasites in immunized mice that may be due to qualitative and/or quantitative difference (small but nonetheless sufficient) in protective cellular populations. In PbATG8-OE-CVac-immunized mice, the liver resident memory T-cell population may have a unique composition to sustain long-term protection. Many studies show a correlation between increased numbers of regional (liver) resident memory T cells (T_RM_) and long-term protection malaria^16,35,66^. Liver-T_RM_ have emerged as a promising target for protecting against malaria as their depletion results in the loss of immunity. Residing in the highly fenestrated blood vessels of the liver (and not in parenchymal tissue), liver-T_RM_ can circulate through the liver to efficiently interact with antigen-presenting cells, (e.g., Kupffer cells, dendritic cells), allowing for the rapid detection of antigen.

In the absence of a role of IFN-γ as a central mediator of protective immune response against malaria in many vaccines, PbATG8-OE-CVac parasites renders its anti-parasitic activity in another way than RAS and other whole sporozoite vaccines. Differences in the effector used by CD8^+^T cells entails that these mutant parasites might interact with the immune system in a unique way that leads to better memory and long-term protection. PbATG8-OE parasites may be a valuable immunological model along with RAS and other GAP vaccines. Deciphering the mechanistic aspects of the host-parasite interaction that leads to differential immune responses will be beneficial to identify novel protective antigens, and thus to develop better vaccines against malaria infections.

## Acknowledgement

The authors are grateful to Fidel Zavala for helpful discussions throughout the course of this study, providing parasite strains and reagents, and critically reading the manuscript. We thank the Insectary and Parasitology Core Facilities of JHMRI, and the technical staff from the Johns Hopkins Microscopy Facility. We also acknowledge Photini Sinnis who provides mCherry-expressing *Plasmodium berghei*. This study was supported by a NIAID R56 AI080631 to IC, a JHMRI pilot grant to IC and the Bloomberg Philanthropies.

## Supplemental information

**Figure S1.**
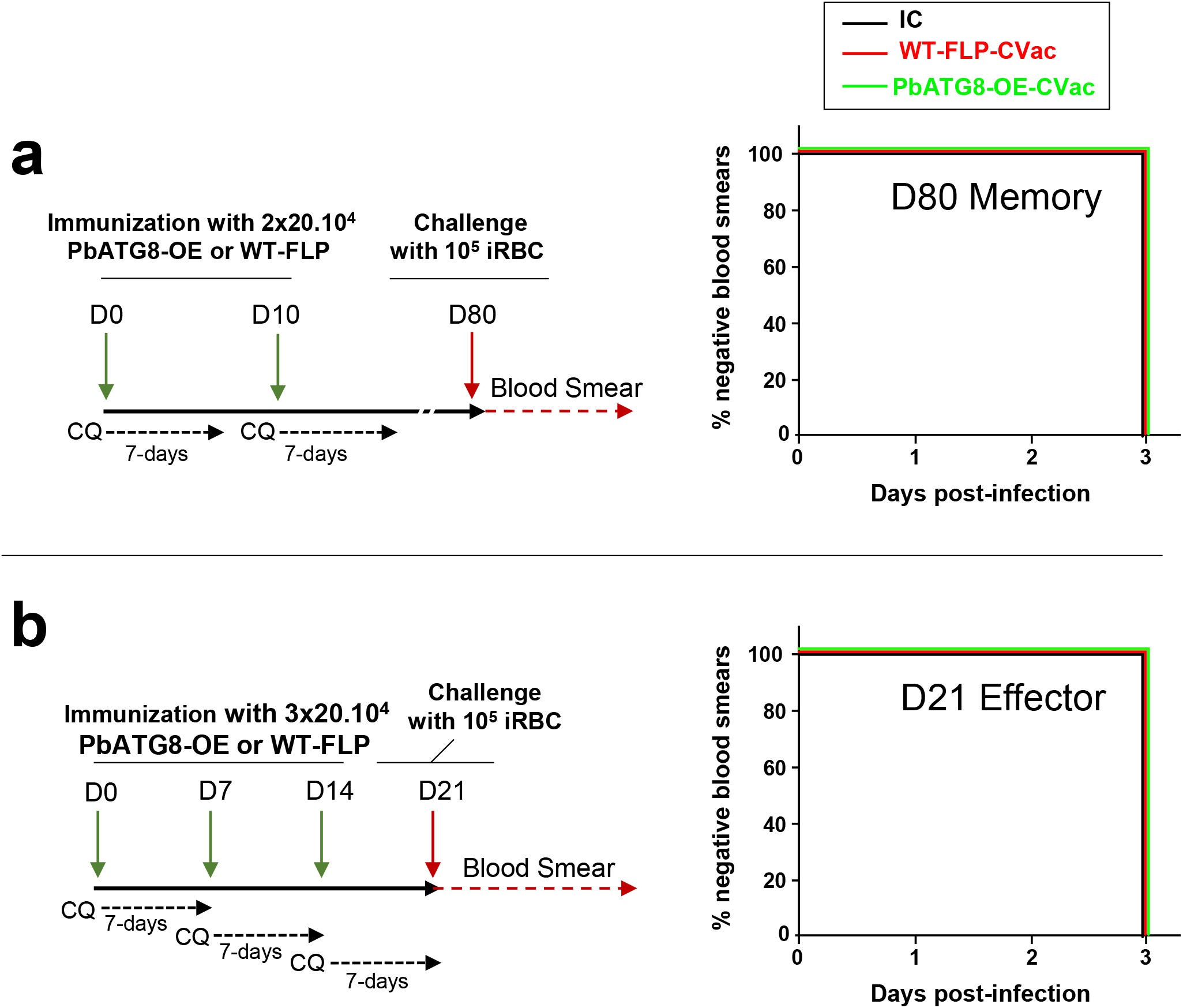
PbATG8-OE-CVac-immunized mice are not protected from blood stage challenge. **a.** Protocol for an immunization with PbATG8-OE-CVac or WT-FLP-CVac parasites and effector challenge with iRBC at D21. Kaplan-Meier survival plots showing days to blood stage infection after challenge. **b.** Protocol for an immunization with PbATG8-OE-CVac or WT-FLP-CVac parasites and memory challenge with iRBC at D80. For a and b, data are representative of one experiment with 5 mice in each group. IC: non-immunized infection control. No statistical significance determined by log-Mantel Cox test.

